# Gastric microbiomes of *Cassiopea* in the Florida Keys are low diversity and *Endozoicomonas-*, *Vibrio-* and *Mycoplasma-dominated*

**DOI:** 10.1101/2024.01.30.577655

**Authors:** Kaden M Muffett, Jessica M Labonté, Maria Pia Miglietta

## Abstract

Interactions with microbial communities fundamentally shape metazoans’ physiology, development, and health across marine ecosystems. This is especially true in zooxanthellate (symbiotic algae-containing) cnidarians. In photosymbiotic anthozoans (eg. shallow water anemones and corals), the key members of the associated microbiota are increasingly well studied, however there is limited data on photosymbiotic scyphozoans (true jellyfish). Using 16S rRNA barcoding, we sampled the internal and external mucus of the zooxanthellate Upside-Down Jellyfish, *Cassiopea xamachana* throughout eight sites covering the full length of the Florida Keys. We find that across sites, these medusae to have low-diversity internal microbial communities distinct from the communities of their external surfaces and their environment.

These internal communities are dominated by only three taxa: *Endozoicomonas* cf. *atrinae*, an uncultured novel *Mycoplasma*, and *Vibrio* cf*. coralliilyticus*. *Cassiopea* bell mucosal samples conform largely to the communities of surrounding sediment with the addition of *Endozoicomonas* cf. *atrinae*. The microbial taxa we identify associated with wild Florida Keys *Cassiopea* bear a strong resemblance to those found within photosymbiotic anthozoans, increasing the known links in ecological position between these groups.

## Introduction

Cnidarian-photosymbiont interactions are central to ecosystem function in coral reefs and beyond. Algal photosymbionts, or zooxanthellae, are present in the well-known stony corals, octocorals, and anemones, but also play important roles in a variety of lesser-known jellyfishes, chief among them, jellyfish of the genus *Cassiopea* (1–4). For symbiotic anthozoans, microbial communities play a key role in maintaining health within this photosymbiont partnership, with community members directly conferring disease resistance and heat tolerance on hosts (5–7).

These microbial communities are often specialized, with distinct corals demonstrating co-evolution with members of their bacterial communities (8), but retain several prevalent genera, such as *Endozoicomonas*, across many hosts (9). While anthozoan communities are under active investigation, few analyses have documented the stability and nature of microbiome communities associated with zooxanthellate jellyfishes, a group with divergent life history traits from Anthozoa but with a comparable obligate symbiosis with dinoflagellates (clade Symbiodiniaceae).

The Upside-Down Jellyfish, *Cassiopea* spp., is a model organism for cnidarian- photosymbiont interactions (10,11). These medusae are residents of tropical and subtropical waters globally and can survive in water that is frequently shallow (as little as 8 cm depth) and hot (up to 33°C for *C. andromeda*) (12,13). *Cassiopea* obligatorily hosts algal symbionts, losing mass even when well fed while in aposymbiotic conditions (14). As in corals and other scyphozoans, mucus plays a central role for *Cassiopea*, presenting a sticky interface between the host and their benthic environment (15,16). In contrast to other symbiotic cnidarians, such as stony corals, *Cassiopea* can tolerate heat and intense temperature fluctuations(17).

Understanding the *Cassiopea* natural microbial community could provide new potential insights into the function and diversity of cnidarian holobionts.

Past jellyfish (class Scyphozoa) microbial studies have focused on the microbiome of lab-reared specimens (18,19) or limited sample sizes (20–22). While these types of studies may be able to track state changes across life stages (18) and between condition types (symbiotic/aposymbiotic) (19), there is little guarantee that the microbial communities of jellyfish kept under prolonged laboratory conditions translate bear resemblance to *in situ* microbiomes. Recent *in situ* studies have provided insights into microbial taxa associated with some scyphozoans ("true jellyfish") but have not included species of the genus *Cassiopea*. In the moon jellyfish, *Aurelia* spp., individuals from the Chinese coast were *Vibrionaceae*-dominated, while samples from the northwestern Atlantic were *Mycoplasma*-dominated(22,23). Between the Indonesian marine lakes Kakaban, Haji Buang, and Tanah Bamban, *Mastigias* microbial community composition varied significantly by lake, but consistently included *Oceanospirillales* (primarily *Endozoicomonas*-like) (20). In samples across taxa, *Mycoplasma* is a common component of the identified gut and tissue microbiome of jellyfishes, though its function remains unknown (22–27). Vertebrate pathogen research is often a focus of scyphozoan microbiome discovery—*Tenacibaculum* has been identified in several scyphozoan species, e.g. *Cotyllorhiza tuberculata, Aurelia aurita,* and *Pelagia noctiluca*, and, as a fish pathogen, which can have important fisheries ramifications (22,23,25,28).

As most studied jellyfish populations occur in coastal waters only sporadically, the relative consistency of *Cassiopea* assemblages provides a system in which location effects can be analyzed at single time points. *Cassiopea* spp. presents an opportunity to act as a comparison group for the microbial communities associated with both other scyphozoans and symbiotic anthozoans. The ubiquity of *Cassiopea* assemblages within the Florida Keys allows for the examination of differences in the microbial community of one population (Florida Keys *Cassiopea*) at one time point across a broad geographic area.

The aim of this work is to identify the core components of the internal and external mucosal microbiomes of *Cassiopea* and compare the microbial communities of *Cassiopea* across sites of the Florida Keys.

## Materials and Methods

### Sampling

A total of 55 medusae were collected by wading in eight sites along the length of the Florida Keys in late August 2021 over eight days (see Table 1 for a list of sites). Species identification for each individual is reported in Muffett & Miglietta 2023 and summarized in Supplementary Table S1 (29). At each site, coordinates, temperature, time of day, depth, salinity, and pH were recorded (Table 1). Salinity was measured with a refractometer, and temperature and pH were measured with an Apera A1209. A minimum of two and a maximum of 10 medusae were sampled per site. At each site and in between each medusa, all equipment was cleaned with absolute ethanol and 10% bleach solution. A 1 L water sample was collected from above (>10 cm above the benthos) each medusa aggregation and filtered (0.2 um pore size, Thermo Scientific Cat No.09-740-30G) with a manual sampler as described in the USGS manual water sampling protocol (30), replicate filters were placed in 15 mL of absolute ethanol or dimethyl sulfoxide ethylenediamine tetraacetic acid saturated salt storage solution (1 L pH 7.5: 93.06 g EDTA, 60 mL 20 % NaOH solution, 20 mL 25 % HCl, 40 mL DMSO, 800 mL water, NaCl to oversaturation; see ref. 30) (commonly referred to as DESS), one filter in each buffer. At each *Cassiopea* medusa aggregation, duplicate scrapings of ∼10 g of sediment collections (substrate sample) were placed in either 15 mL of absolute ethanol or DESS storage solution.

**Table 1:**
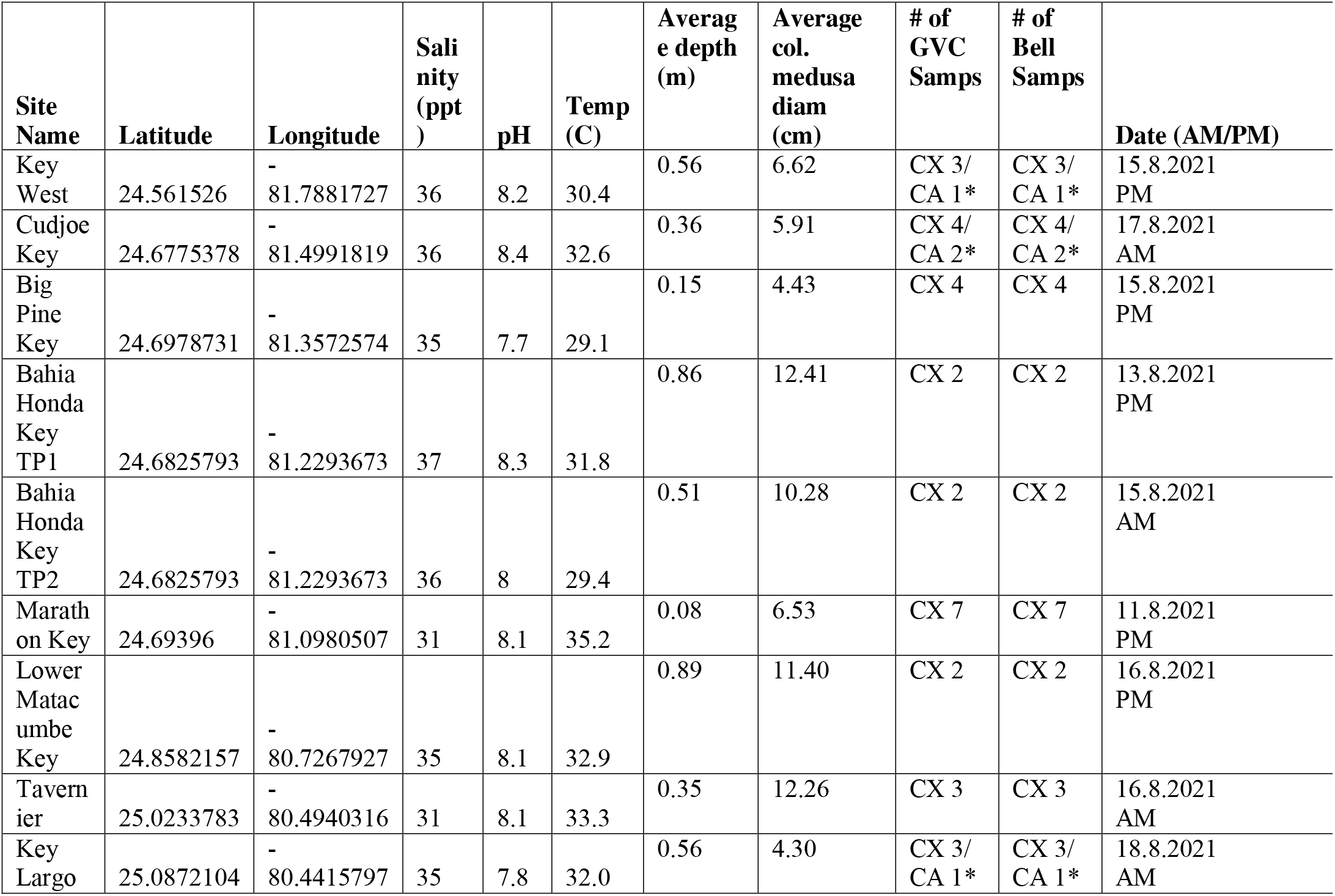
List of collection sites. Table with latitude, longitude, salinity, pH, surface temperature, average diameter of collected medusae from the site, and date of collection. Number of paired GVC and bell swab samples per collection are for *Cassiopea xamachana* (CX). For collections that included *C. andromeda* mitotyped medusae, the number of *C. andromeda* samples is presented after the dash (CA). These values only include samples collected in DESS.

Each *Cassiopea* medusa was collected from shallow waters <1.5 m within a single assemblage at each site. Medusae were gently grasped by the oral arms, lifted from the water, swabbed along the apex of the bell, then placed bell-down onto a pre-prepared dissection plate (Figure 1). External bell swabs were done in replicate and placed in 3 mL DESS or absolute ethanol. Medusa oral arm base was bisected between oral arm groups, and replicate swabs were inserted and circled the digestive cavity. One internal swab was placed in DESS, the other in ethanol, as with external swabs. A section of bell tissue was then collected and placed in ethanol for mitochondrial species confirmation (see 28). The size of each medusa was measured, and individual medusae were photographed. As sampling was done in a nonsterile environment, blanks for buffer with swab and buffer with filter were also taken.

**Figure 1:**
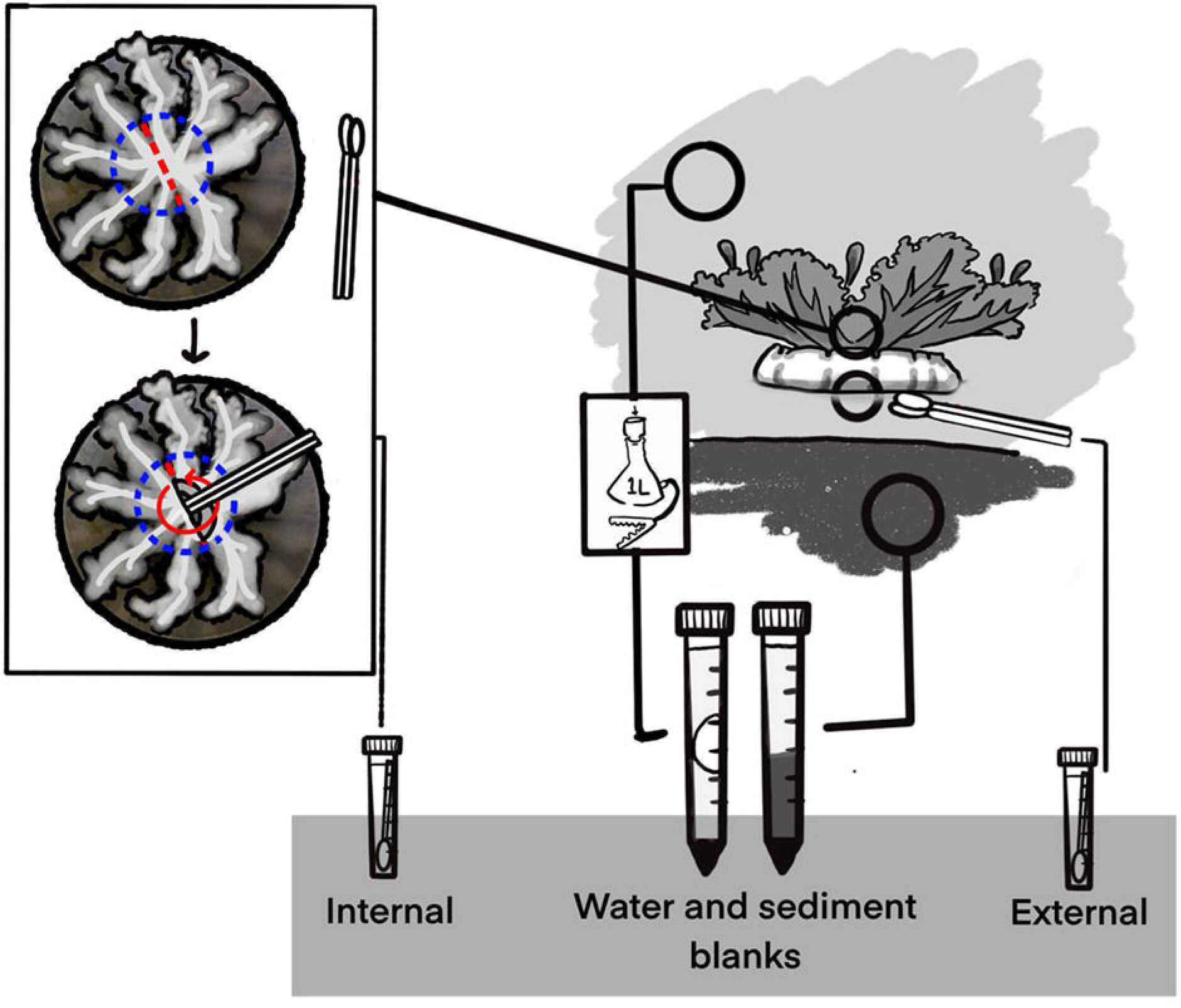
Sampling strategy for medusae collection onsite. Demonstration of the typical orientation of medusa (right upper) and swab locations (apical bell surface and internal to medusa). GVC access strategy is displayed on the left and relevant environmental samples (water filter, substrate sample) are displayed on the right.

### DNA Extraction and Sequencing

Samples in DESS and EtOH were maintained at room temperature for three to ten days and then frozen. DESS can be stored at room temperature for at least thirty days without significantly compromising community integrity (31). DNA from all samples (see Supplementary Table S1) was then extracted using Zymobiomics DNA Kits (PN# D4300T). All samples were pre-tested through the ampliciation of the 16S rRNA gene (variable region V4) amplifications. Only samples that produced visible bands for the V4 region were sent for 16S rRNA V3-V4 preparation and sequencing. While DESS preserved samples yielded good amplification results, EtOH-preserved samples largely failed to produce adequate amplification. As such, only DESS-stored samples were analyzed. Sequencing was performed by the University of Texas Medical Branch Sequencing Lab. Library was prepared with Zymo Quick 16S V3-V4 kit (PN #D6400) in 20 uL reactions (10 uL Quick-16S qPCR premix, 4 uL Quick-16S V3-V4 primers (341f: (CCTACGGGDGGCWGCAG, CCTAYGGGGYGCWGCAG) and 806r:

(GACTACNVGGGTMTCTAATCC)), 4 uL water and 2 uL sample). qPCR was run using Roche LightCycler 480 Instrument II (10min 95C, [30sec 95C, 30sec 55C, 3min 72C]x20). PCRs were enzymatically cleaned according to Zymo Quick-16S kit protocols, and barcodes were added in a secondary reaction at 20 uL (10 min 95°C, [30 sec 95°C, 30 sec 55°C, 3 min 72°C]x5). Plates included two replicate microbial community standards (Zymobiomics Cat. No. D6300) and two PCR negative controls. Samples were loaded at 10 pM, with 15% 10 pM PhiX sequencing control, and run on Illumina MiSeq with a 600 cycle v3 kit.

### Data preprocessing

Data generated was run through the DADA2 v 1.26 pipeline on R v 4.4 with only limited deviations from the standard protocols for ASV generation(32,33). Briefly, sequences were filtered and trimmed (“filterAndTrim(truncLen=c(285,230), maxN=0, maxEE=c(4,4), truncQ=2,trimLeft = c(17,21)”), errors were error corrected (“learnErrors”, “dada”), and sequences were merged (“mergePairs”). Reads between 380 and 458 bp were retained. Chimeric reads were removed (“removeBimeraDenovo”), then taxonomy was assigned using the Silva v 138.1 database (“assignTaxonomy”). Retention metrics were retained and these metrics along with the full code can be found on the github (https://github.com/kadenmuffett/Cassiopea-Code-Microbiome-Repository). Sequence and metadata tables were imported into phyloseq v 1.42, where reads assigned to mitochondria and chloroplast were removed. Contaminants were removed using the decontam package v. 1.18, identifying and removing 47 ASVs based on differential abundance in swab and buffer controls(34,35).

For all downstream analyses, dataset was limited to those stored in DESS and medusae without a *C. andromeda* mitotype. This includes 60 paired bell and GVC samples (30 of each), 8 water samples and 8 substrate samples with 6.47 mil sequences across 34808 taxa.The complete dataset (NCBI Bioproject Accession number PRJNA1020388) contains 12.78 million reads.

### Statistical analyses

Refraction curves were generated in vegan 2.6.6.1 (“rarecurve”) (SFIG. 1)(36). Shannon diversity was computed with vegan and observed diversity was ploted with phyloseq (“estimate_richness”, “plot_richness”)(34). Faith’s phylogenetic diversity was computed using the MiscMetabar v 0.10.1, phanghorn v 2.12.1, metagMisc v 0.5.0 and PhyloseqMeasures v 2.1 packages by first generating an ML tree from all sequences over 300 reads across the dataset, then computing per sample Faith’s diversity (37–42). Faith’s diversity was displayed with ggstatsplot v 0.13.0 (43). Bray curtis and Euclidean distance were computed in phyloseq with “distance” to “ordinate” to “plot_ordination”. Between group comparisons were completed with the pairwiseAdonis package v 0.4.1 (“ pairwise.adonis2”) on Euclidean distance measurements (44). Statistical comparisons were computed with Kruskal-Wallis using the Kruskal.test and pairwise comparisons using the “kruskal.test” and “dunn.test” commands with holm p adjustment (45).

Data transmutation for starplots and display was assisted by the speedyseq package v 0.5.3.9021 (46). Star plots were made using the stars package v 0.6-7 (47). Differential abundance was calculated with ALDEx2 v 1.38.0 following established pipelines for microbiome data (48,49). Dirichlet Multinomial mixture analysis, as described by Holmes and implemented in the DirichletMultinomial package v 1.48 (50,51). Core microbiome was computed with the microbiome package v 1.28.0(52).

### Ethics Statement

This work was performed on lower level invertebrates not subject to Animal Use Protocols or within the purview of Texas A&M University IRB approval. Permitting for these specimens was waived by Florida Fish and Wildlife Conservation Commision.

## Results

### Data description

We analyzed the microbiome associated with 34 medusae from eight sites within the Florida Keys. For each medusa, we analyzed the microbial composition of their gastrovascular cavity (GVC) and external or bell mucus (herein referred to as “bell”), as well as their substrate (Sub) or water environment (WTR). All medusae were identified as *Cassiopea xamachana,* with the exception of four medusae identified as mitochondrial haplotype *C. andromeda* found in Key West, Cudjoe Key, and Key Largo (see Supplementary Table S1). 10.13 and 9.07 million reads were retained after filtering (“filterAndTrim”) and merging (“mergePairs”) respectively (for all retention statistics, see supplementary table S2) . There were 8.91 million sequences after chimera removal. After primary phyloseq filtration, the dataset included 6.47 million sequences retained in 34808 unique ASVs across 76 total samples (30 bell, 30 GVC, 8 water, 8 substrate). Reference sequences for top ASVs are also available in the supplementary material (table S3).

### Distribution of sequences in full dataset

The dataset generated was predominantly bacteria with few archael reads. At the phylum level, 57% of sequences were Proteobacteria, 7% were Firmicutes, 13% were Bacteroidota, and 4% were Cyanobacteria (Fig 2). No other phylum had greater than 3% relative abundance in the dataset.

**Figure 2.**
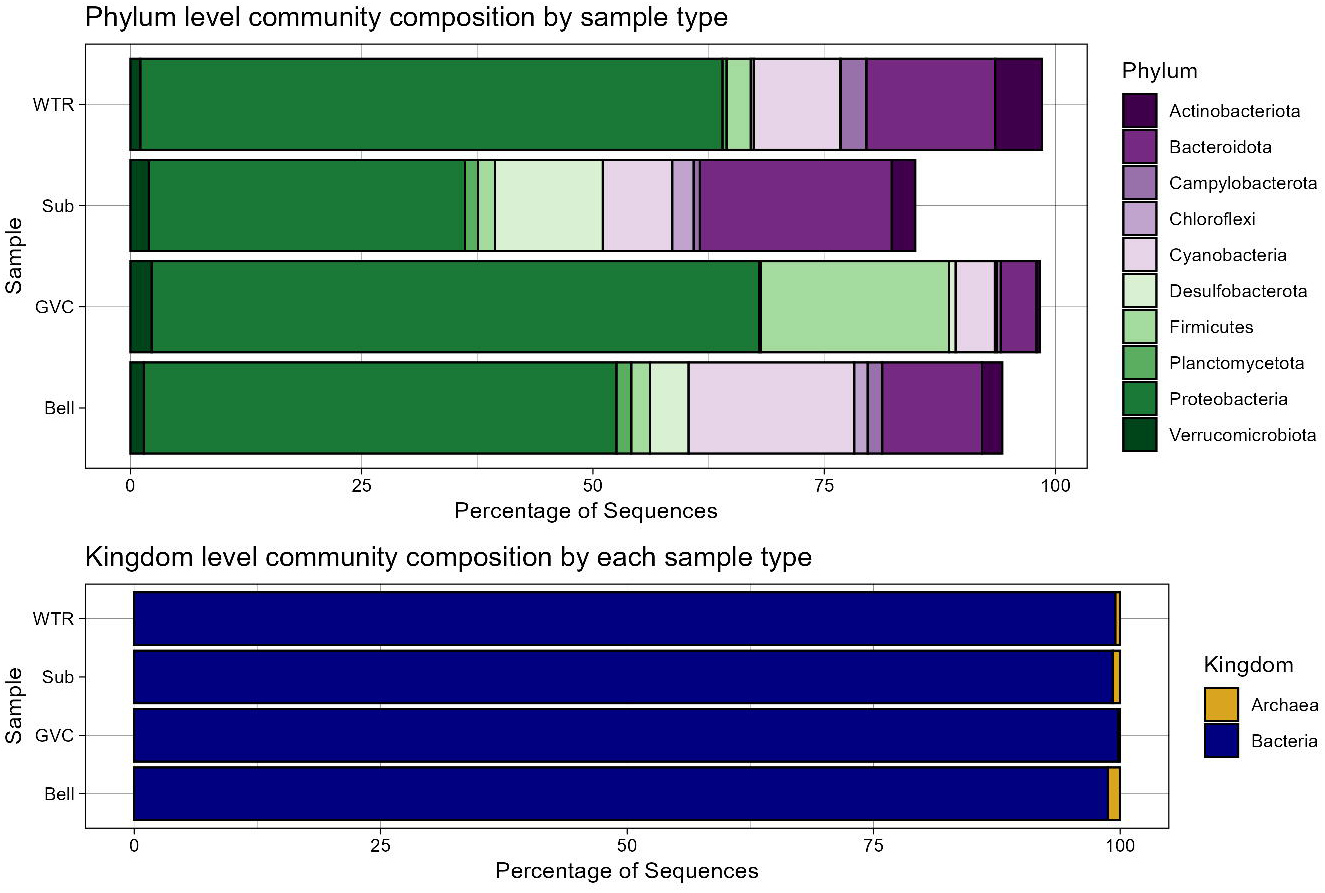
Top ten (top) phyla and (bottom) class relative abundances of sequences in samples aggregated by sample type.

At the class level, Gammaproteobacteria comprised 41% of all sequences.

Alphaproteobacteria, the next most abundant, comprised 17%. Bacteriodia (12%), Bacilli (6%), Cyanobacteriia (4%), and Desulfobacteria (3%), were the other classes with a relative abundance above 2%. No Archaean group was prevalent above a relative abundance of 0.25%, with the three most abundant classes being Halobacterota (0.25%), Thermoplasmatota (0.21%) and Nanoarchaeota (0.22%).

### Distribution of sequences within gastrovascular cavity (GVC) samples

Within gastrovascular cavity samples, 75% of all sequences belonged to just three orders, Pseudomonadales (39%), Mycoplasmatales (24%), and Enterobacterales (13%). The majority of Pseudomonadales sequences identified within the GVC were dispersed across very few OTUs, with 66% identified as *Endozoicomonas atrinae* ASV1 and 8% as *E. atrinae* ASV4. This was true of the Mycoplasmatales as well, with 77% of associated gut sequences identified as *Mycoplasma* sp. ASV2 or *Mycoplasma* sp. ASV9 (15%).

### Distribution of sequences within bell mucus samples

At the phylum level, within the bell mucosal swabs, 55% of the sequences were Proteobacteria, 14.5% Bacteroidota, 5.1% Cyanobacteria, and 6% Desulfobacterota.

At the order level, the most abundant groups were Pseudomonadales (23%), Chromatiales (6%), Rhodobacterales (7%), Desulfobacterales (5%), and Flavobacteriales (5%). *Endozoicomonas atrinae* ASV1 was the single most abundant surface mucus ASV at a mean of 12%. The next most abundant were Synechococcus CC9902 ASV15 (1.3%), *Endozoicomonas atrinae* ASV4 (1.1%) and Chromatiales *Candidatus Thiobios* ASV6 (0.8%).

### Substrate and water samples

Water samples shared some ASVs with the bell mucus of medusae, the largest contributing ASV was Chromatiales *Candidatus Thiobios* sp. ASV6 (8%). Additional prominent taxa were Rhodobacteraceae *HIMB11* ASV12 (6%), *HIMB11* ASV19 (5%), and SAR11 Clade III ASV 10 (4%). By contrast, substrate samples had no ASVs surpassing a mean of 1% abundance across sites, with the highest two ASVs Sandaracinaceae ASV94 and *Actibacter* ASV105 reaching 0.6% mean abundance across sediment samples (see table 2; extended table is table S4).

**Table 2:** Top 5 ASVs for each sample type. Table with ASVs ordered from highest to lowest average proportion in samples of each type dispayed as a percentage of composition within that sample type. Includes Silva 138 taxanomic assignment for each ASV.

|  | 1 | 2 | 3 | 4 | 5 |
| --- | --- | --- | --- | --- | --- |
|  | ASV1<br>*Endozoicomonas atrinae* | ASV15<br>*Synechococcus*<br>CC9902 | ASV4<br>*Endozoicomonas atrinae* | ASV6 *Candidatus Thiobios* | ASV10 SAR11<br>Clade III |
| Bell | 11.585 % | 1.272 % | 1.105 % | 0.832 % | 0.709 % |
|  | ASV94<br>Sandaracinaceae | ASV105<br>*Actibacter* | ASV95<br>Desulfocapsaceae | ASV169<br>*Robiginitalea* | ASV358<br>*Clostridiisalibacter* |
| Sub | 0.625 % | 0.605% | 0.566% | 0.492% | 0.474% |
|  | ASV6 *Candidatus Thiobios* | ASV12<br>Rhodobacteraceae<br>*HIMB11* | ASV19<br>Rhodobacteraceae<br>*HIMB11* | ASV10 SAR11<br>Clade III | ASV18<br>*Marinobacterium* |
| WTR | 7.521 % | 5.839% | 4.725% | 4.497% | 2.711% |
|  | ASV1<br>*Endozoicomonas atrinae* | ASV2<br>*Mycoplasma* | ASV9 *Mycoplasma* | ASV4<br>*Endozoicomonas atrinae* | ASV11 *Vibrio coralliilyticus* |
| GVC | 34.133% | 20.099% | 2.823% | 2.327% | 2.094% |

### Diversity

Shannon diversity varied by sample type (Kruskal-Wallis: chi-squared = 48.8, df = 3, p-value =1.412e-10). Shannon diversity had a mean of 2.30 in GVCs, 4.01 in water samples, 5.63 in substrate samples and 6.56 in bell samples. Shannon diversity indices for the gastrovascular samples varied from substrate samples and bell samples (Kruskal-Wallis Dunn test with Holm correction: GVC-Bell, p-val=0.00; GVC-Sub, p-val= 0.00; GVC-Water, p-val=0.07; Bell-Sub, p- val=0.07; Bell-Water, p-val=0.06; Water-Sub, p-val=0.01). Shannon diversity was not explained by location (Kruskal-Wallis: chi-squared = 9.7, df = 7, p-value =0.21)(Fig. 3).

**Figure 3.**
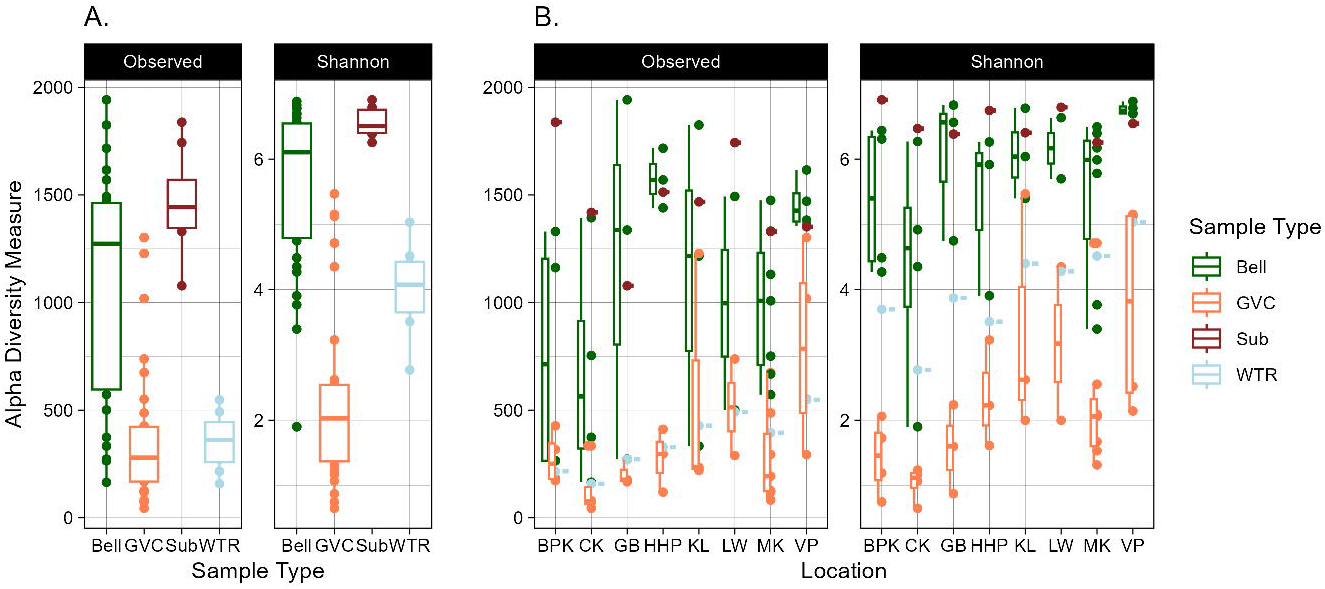
Shannon diversity (a) by sample type and (b + c) location. GVC, bell, substrate and water Shannon diversity are shown across both the entire dataset or when segmented by sample type across sites with (b) GVC and (c) bell alpha diversity across sites. All p values of holm-adj Dunn comparison of means tests below a = 0.05 are displayed, with statistical test details displayed above each plot.

**Figure 4.**
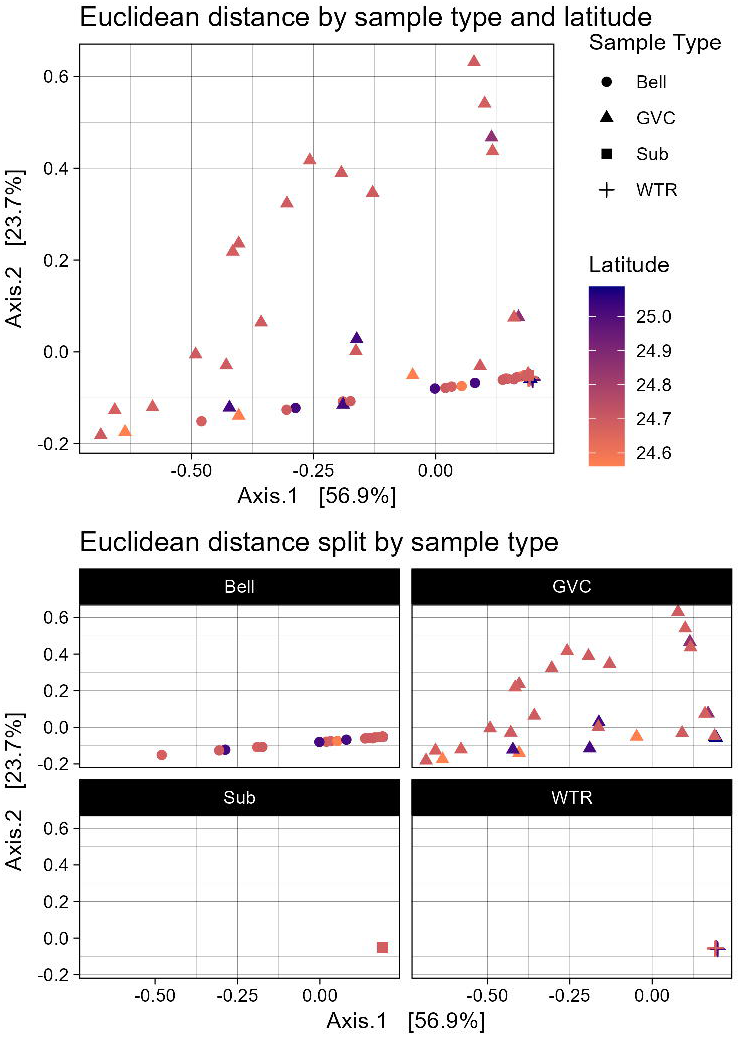
PCA of Euclidean distance of samples. PCA is colored by sample collection latitude from low (orange) to high (purple). Sample type is designated by shape, circle (Bell), triangle (GVC), square (Substrate) and cross (Water). The plots show the same PCA faceted by sample type.

### Beta diversity

Euclidean distance showed differences between medusa and non-medusa samples (pairwise.adonis, GVC vs bell: p=0.001, bell vs sub: p=0.049, bell vs water: p=0.022, GVC vs sub: p=0.001, GVC vs water: p=0.001, sub v water p=0.001). Variability along the primary axis was found in both GVC and bell samples, while variability along the secondary axis was found mainly in GVC samples. The location of collection had no clear impact on beta diversity.

Faith’s diversity of communities were distinct from each other (Kruskal-Wallis test, chi- squared = 44.1, df = 3, p-value = 1.4e-09). GVC communities were distinct from bell and substrate comparison groups, while bells were distinct from water and gastrovascular cavities (Dunn test with holm correction) (Fig. 5).

**Figure 5.**
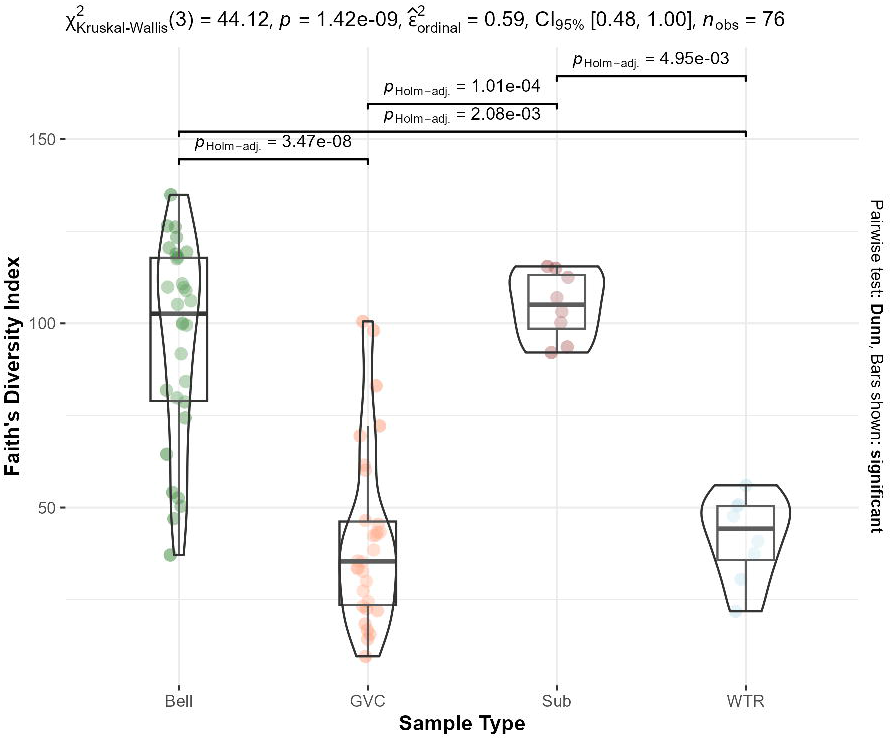
Violin plot of Faith’s diversity. Samples are colored by sample type. Kruskal-Wallis chi-sq, pvalue and pairwise dunn test with holm correction p-values are displayed. Differences between GVC-water and substrate-bell are not significant.

### Core microbiome

Core microbiome for the purposes of this study was evaluated based on 70% occurrence of an ASV and a relative abundance of > 0.1% of reads in all 34 gastrovascular cavity samples or external bell samples. This is in line with thresholds for other core microbiome work (53). The most abundant ASV, *Endozoicomonas* ASV1, was present in 90% of GVC samples at a relative abundance of > 0.1% of reads, ASV2 and ASV4 were present in 80% . Using less stringent criteria (>70% of samples, any abundance), ASV 11, ASV 132 and ASV 6 become part of the core microbiome (Fig 6). These ASVs are across only three genera- *Endozoicommonas*, *Mycoplasma* and *Vibrio* (full taxonomic details of ASVs can be found in table S5).

**Figure 6.**
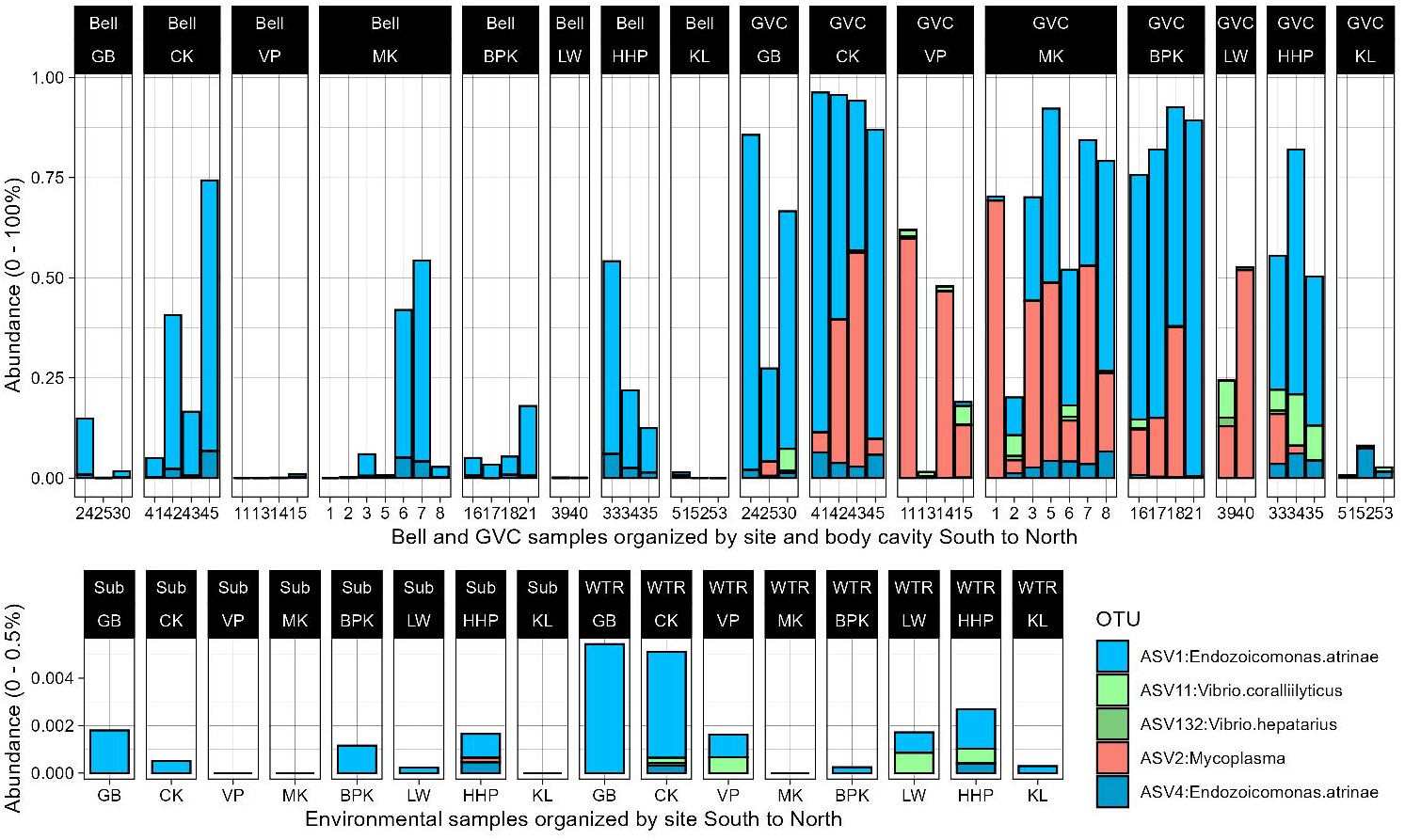
Abundance of Cassiopea core GVC microbiome across dataset. The top five ASVs from the gut core microbiome across all sample types (Bell, GVC, Substrate (Sub) and Water (WTR)) organized from furthest south to furthest north sites. Note the difference in y axis maxima between bell and GVC samples (top) and substrate and water samples (bottom).

In bell samples, ASV1, ASV104, and ASV86 have 73%, 70% and 70% relative abundance in samples at a 0.1% threshold. ASV6, ASV95, ASV37, ASV105, and ASV135 are present at any level in >70% of bell samples. These ASVs cover the genera *Endozoicommonas*, *Candidatus Thiobios*, *Methyloceanibacter, Ruegeria* and unknown genera from Desulfocapsaceae and Actibacter (Fig. 7).

**Figure 7.**
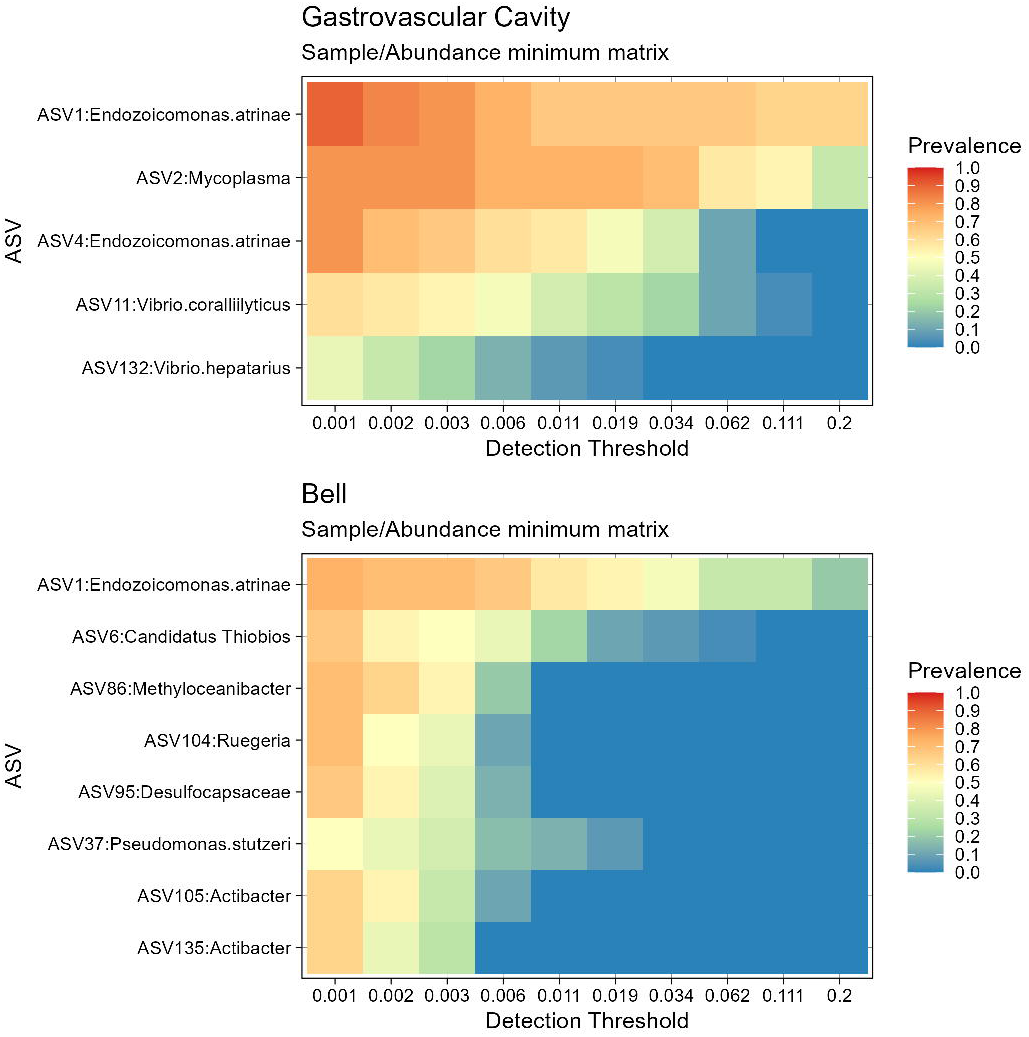
Core microbiome prevalence in GVC and bell. Heatmap of gastrovascular cavity (top) and bell (bottom) shaded by prevalence (0-1 scale) vs detection threshold (proportions of reads in samples) across the ASVs identified as core in each group.

In terms of abundance, internal core microbiome components are not just present, but highly dominant in *Cassiopea*’s gut. Some of these taxa are highly restricted to GVC samples (*Mycoplasma*), while others are prevalent across body surfaces (*Endozoicomonas*) (Fig. 7).

The vast majority of ASVs were not differentially abundant between bell and GVC samples. Differential abundance (Aldex2, min effect size absolute value >[±2]) between the gastric and bell surfaces is restricted to the novel *Mycoplasma* group (ASV2), which is completely lacking in external samples (Fig 8). No features were differentially abundant between substrate and bell samples (Sfig 2).

**Figure 8.**
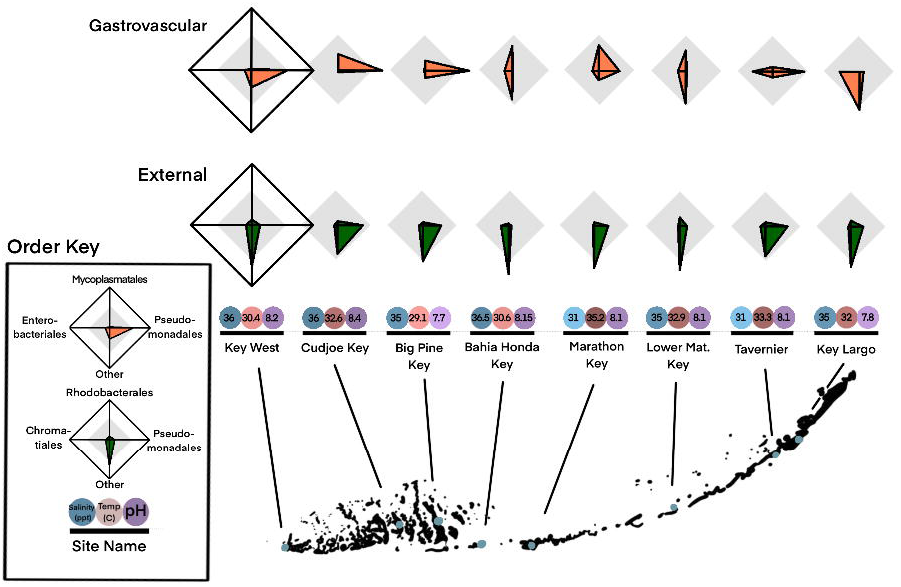
Aldex2 plot of effect size of all ASVs between bell and GVC samples. Only ASV2 has an effect >[± 2].

Dirichlet multinomial mixture (DMM) analysis identified specific proportions of many of these core microbiome as valuable for category differentiation. Within animal-associated samples, model probability value was optimal for two communities, one GVC and one bell without subcommunities. False assignment rate (mismatch between predicted community and true sample type) was present in 5/60 samples. ASV1 amount was a core group differentiator, with fewer meaningful ASVs defining the GVC community than the bell community (Fig. 9).

**Figure 9.**
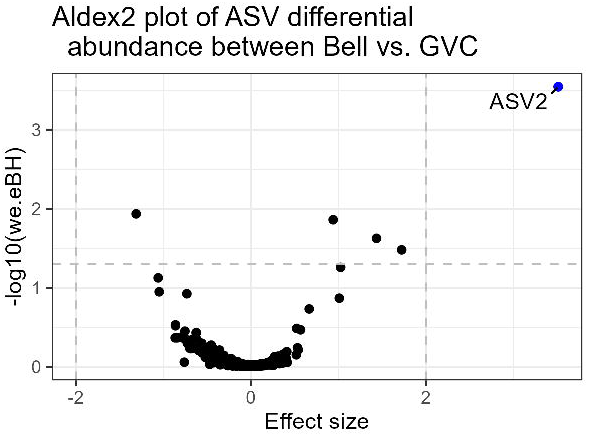
Most impactful ASVs for Dirichlet multinomial modeling. Top 0.5% most impactful drivers of body cavity prediction in GVC (top) and bell (bottom) Dirichlet multinomial models ordered from largest to smallest effect size.

Despite DMM identifying all gastric samples as belonging to the same community, relative proportions of those community members differed dramatically, varying substantially in abundance of *Endozoicommonas*, *Mycoplasma* and *Vibrio*. Variation within bell samples primarily occurred with proportion of *Endozoicommonas,* but was not reliably distinguishable from substrate samples by DMM (Fig. 10).

**Figure 10.**
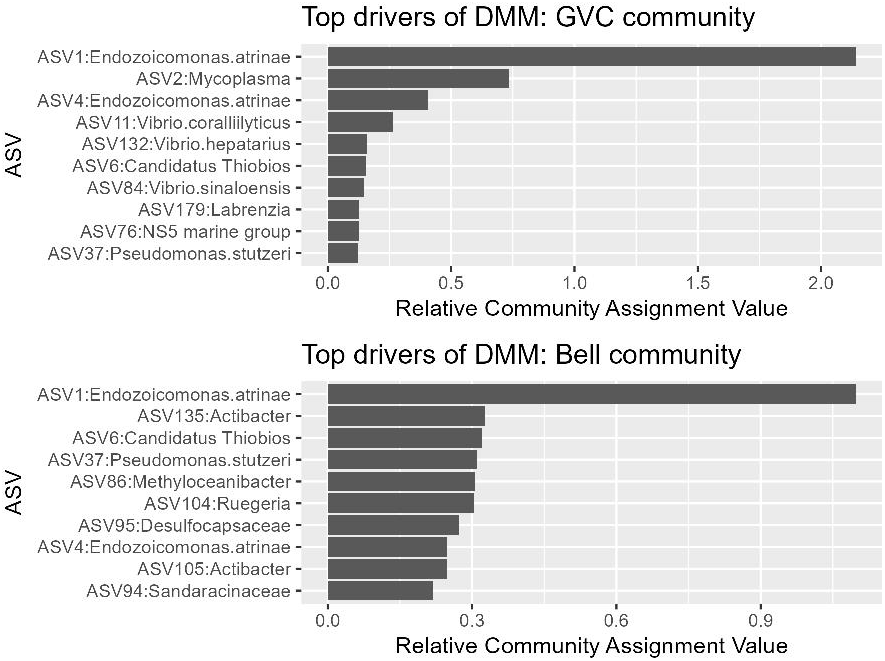
Star plots of community composition across sites. Star plots for internal (GVC), external (Bell) mucus swabs, water (WTR) and substrate (Sub) sample averages at each site separated by order (three most prominent orders and "other"). Each spoke of the star plot ranges from 0% (center) to 100%. **Internal** group, starting from the 3’ position and going counterclockwise: Pseudomonadales, Mycoplasmatales, Enterobacteriales and “Other”. **External** group spokes (going counterclockwise): Pseudomonadales, Rhodobacterales, Chromatiales, and “Other”. **Water** group spokes (going counterclockwise): Pseudomonadales, Rhodobacterales, Flavobacteriales, and “Other”. **Substrate** group spokes (going counterclockwise): Rhodobacterales, Flavobacteriales, Desulfobacterales, and “Other”. Each site’s measured environmental conditions (salinity, water surface temperature, and pH) are presented below star plots, colored by intensity (low to high represented by pale to dark) . Sites are mapped onto their location within the Florida Keys.

#### Vibrio

ASVs identical to *Vibrio coralliilyticus* were found across *Cassiopea* gastric samples, with abundance of up to 10% of reads within the sample. This far surpasses rates in the surrounding water and substrate samples (ASV11 averages: Bell: 0.05%, WTR: 0.03%, Sub: 0%, GVC: 2.09%) (Fig. 11).

**Figure 11.**
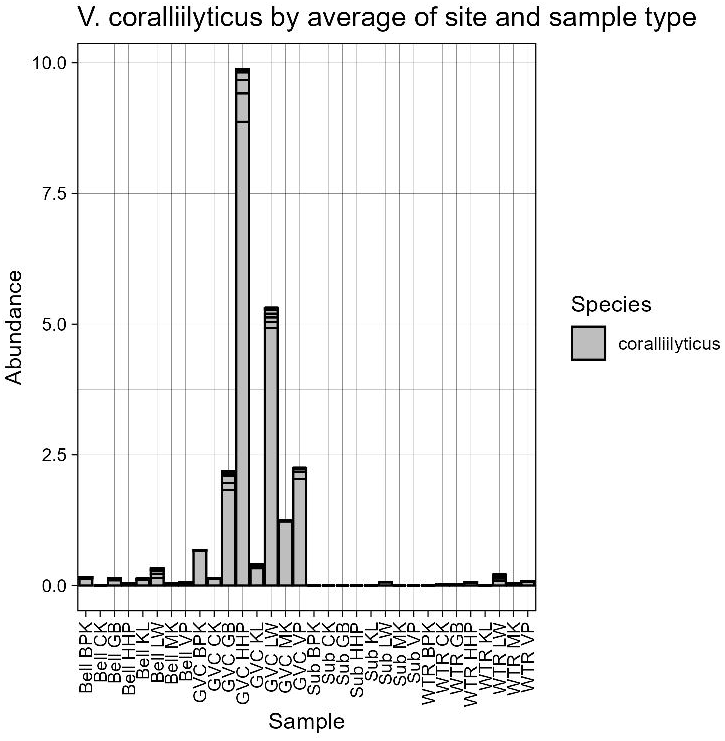
Percent of reads (0%-10%) assigned to *V. coralliilyticus* ASVs in bell, GVC, substrate and water samples averaged by site. Organized by body cavity.

### Host medusa species identity

The mitochondrial haplotype of the non-native *Cassiopea*, *Cassiopea andromeda,* a cryptic sister species to *C. xamachana*, was found in four of the 34 medusae sampled. The four *C. andromeda* mitotype individuals included were from the Key Largo (1), Cudjoe Key (2), and Key West (1) sites. While these were excluded from the primary analysis, *Cassiopea andromeda* showed no divergence from the trends of the *C. xamachana* simultaneously collected (Fig. 12).

**Figure 12.**
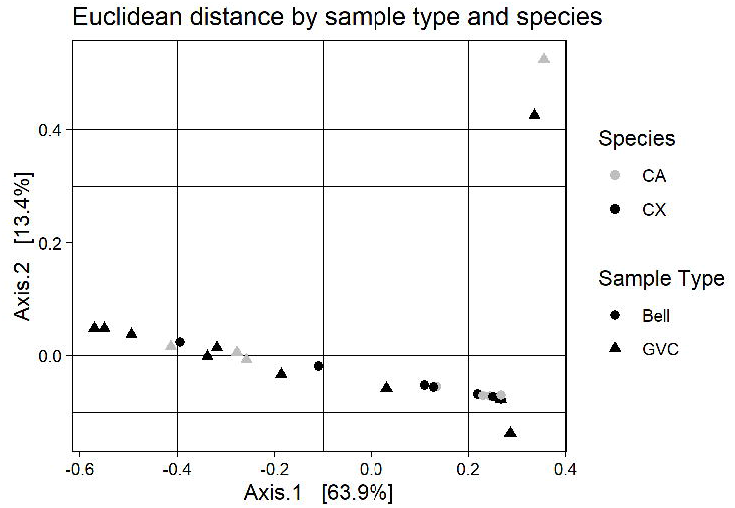
PCA of Euclidean distance between *Cassiopea* species. *Cassiopea andromeda* (CA: grey) and *Cassiopea xamachana* (CX: black) samples at the sites were they cooccurred (Cudjoe Key, Garrison Bight and Key Largo).

## Discussion

*Cassiopea* medusae show consistent differences in the microbial communities present on their bell and gastric surfaces. Gastrovascular cavity samples have lower diversity than the sediment, external surfaces, and host a novel *Mycoplasma* lineage. This low diversity is consistent with results for multiple other scyphozoans— including *Cotylorhiza tuberculata* (25). *Cassiopea* also appears to have some variation in community composition between sites, a feature consistent with results from other species, such as *Aurelia, Mastigias,* and *Tipedalia* (20,24).

### A bifurcated outer and inner microbiome

Across all sites, the gastrovascular cavities of *Cassiopea* medusae showed a core microbiome comprised of taxa previously found in photosymbiotic cnidarians. The external microbial community of *Cassiopea* from the Florida Keys adheres fairly closely to the epibenthic microbial community at each site, with the addition of *Endozoicomonas*. ASV1 *Endozoicomonas* cf. *atrinae* may be a primary mucosal component and extend to the exterior of the body through mucus exchange, as it is less prevalent on the bell surface than internal for most individuals.

Alternitavely, ASV1 may be a common mucosal resident with no function for the host.

The abundant *Mycoplasma* and *Vibrio* cf. *coralliilyticus* from the gastric cavity of *Cassiopea* were not found in any external mucus samples and found at far lower abundances in external mucosal samples respectively.

The high diversity of the *Cassiopea* bell microbiome found in this study may be unique to *Cassiopea*. Previous work has reported in the low-diversity mucosa, oral arms and umbrella of a previously studied pelagic rhizostome, *Rhizostoma pulmo,* a likely result of the constant contact of *Cassiopea* with the epibenthos (26). The limited overlap between diverse bell communities and low diversity gut communities in *Cassiopea* is notable and consistent with the significant differences in body compartment microbiome of other scyphozoans and anthozoans (21,24,54). While the lack of diversity in scyphozoan microbiomes may seem distinct from many described stony coral microbial communities, these stony coral microbial profiles are often produced using whole body sampling. Lower diversity gastrovascular cavities with distinct community composition may also be a feature of stony corals (55–57).

Microbial communities across sites within this study were not significantly different from each other, as gastrovascular cavity components remained dominated by three taxa throughout the Keys. This microbiome bears limited resemblance to those sampled in common garden experiments and no resemblance to lab-held lines, demonstrating that “core” here should not be misconstrued as obligate (19,58). The lack of obligate bacterial taxa in the *Cassiopea* gastrovascular cavity is also consistent with other microbial hosts, notably the anemone *Exaiptasia diaphana,* another successful model cnidarian (1). While *Cassiopea* require their algal symbiont, bacterial partners may be transient, and bacteria may have far less significance to host health for *Cassiopea* than for specialized obligate hosts (e.g., leafhoppers) (59–62). Without a better understanding of specific metabolic function of different bacterial associates, microbiome importance cannot be determined. We identified *Endozoicomonas*, *Mycoplasma,* and *Vibrio* as the three most common bacterial genera found within the gastrovascular cavity of *Cassiopea*. These are all commonly associated with coral microbial communities (8,9,63). Below, we provide information on these common bacterial components of cnidarian holobionts.

### *Endozoicomonas* functions in microbiomes

The large proportion of *Endozoicomonas* in internal and external samples fits within a broader framework of both *Endozoicomonas* and known cnidarian-associated microbial communities; *Endozoicomonas* strains are present in a variety of invertebrates, including many corals, sponges, sea anemones, scyphozoans and cubozoans (9,63–66). Some hosts house multiple distinct *Endozoicomonas* genotypes, suggesting that different *Endozoicomonas* clades may provide different metabolic services to a host (e.g., Vitamin B12 metabolism) (67).

*Endozoicomonas* is more commonly identified in healthy scleractinian corals than bleached and diseased ones (68). For these corals, it may play a role in a variety of fundamental *Symbiodinium*–host interactions, including nutrient transport and symbiont cell breakdown (67). Notably, *Endozoicomonas* is not a consistent marker of microbiome health, its disappearance and reappearance without apparent host health declines has been identified in octocorals (2,66). In some coral, members of *Endozoicomonas* are found in close proximity to *Symbiodinium,* it is unknown whether this feature is shared with *Cassiopea* (67).

Within scyphozoa, *Endozoicomonas* is abundant in the microbial community of the rhizostome scyphozoan *Mastigias* (20). While *Mastigias*, like *Cassiopea*, host zooxanthellae from the family *Symbiodiniaceae*, not all symbiotic rhizostomes harbor the microbe, and not all taxa with the microbe are symbiotic (20,25). *Endozoicomonas* may be common across medusae, but the difficulty of culturing some strains may have excluded them from the results of past culture-based analyses (66,67). Greater clarity is needed on the function of *Endozoicomonas* for these medusae and other taxa.

### Mycoplasmataceae: a poorly understood cnidarian microbiome resident

*Mycoplasma*, like *Endozoicomonas*, is present in studied coral and scyphozoan microbiomes (9,22,23,25). While intracellular parasitism is common for members of Mycoplasmataceae, those associated with cnidarians may be located extracellularly and exist as opportunistic commensals (2,69). *Mycoplasma* sp. is noted as a potential endosymbiont in lab-raised *Aurelia* polyps, while adults from many jellyfish populations are known to have them (22,24,27). The high divergence between this *Mycoplasma* and others of the genus (closest sequence identity on NCBI at 89%: Accn. No. NR_115937) suggests that this may be a hitherto unknown group. Whether this *Mycoplasma* represents an endosymbiont, a result of prey capture, or a parasite, its exclusivity to the gastrovascular tract in *Cassiopea* sampled within the Keys is notable, as other studies have not demonstrated gastrovascular cavity localization (24,27).

### Vibrionaceae

Vibrionaceae is both a core part of coral microbiomes (9,70) and known to facilitate metamorphosis within *Cassiopea* (71,72), however, it is a highly diverse family. While *Vibrio* spp. are found in gut communities of presumed healthy scyphozoans (Daley et al. 2016, Peng et al. 2021), they can also be associated with senescence and disease (Kramar et al., 2019; Tinta et al., 2012). For *Cassiopea* specifically, an increase in *Vibrio* has previously been associated with an aposymbiotic or stressed state (58). Given the variation in Vibrionaceae species and strains, family-wide statements on the beneficial or harmful nature of Vibrionaceae within *Cassiopea* gastrovascular cavities should not be made.

#### Vibrio coralliilyticus

In this dataset we found ample signal of a 16S V3-V4 with 100% sequence match to multiple *Vibrio* strains, including *V. coralliilyticus* and the oyster pathogen, *V. tubiashii*. As *Vibrio* species are highly varied, the sequence match does not mean assured pathogenicity towards any taxa (75–77). The apparent health of the medusae from whom these samples were collected suggest a either a lack of impact of this specific group on the host or a host-microbe facultative interaction between this *Vibrio* and *Cassiopea.* While there is data on *Cassiopea* microbiome antimicrobial isolates and *Cassiopea* mortality when faced with *Serratia marcescens* infection, we lack adequate details on the *Vibrio* – *Cassiopea* system to determine whether this represents a lowlying widespread infection or a mutualism (78,79).

### Drivers of external microbiome

While the external microbiome was not distinct from the sediment around it, it is worth discussing the major groups more common externally than internally. While the core bell microbiome represents a substrate-like community, the bell core microbiome includes representatives from families and genera with a variety of potential metabolic capacities. The most plentiful of these ASVs was *Candidatus Thiobios*, known as a sulfur-oxidizing epibiont of the *Zoothamnium niveum* (80,81). As this bacteria can be found in the Florida Keys, their appearance on *Cassiopea* may be circumstantial, but is consistent across medusae (80). Isolates from *Methyloceanibacter* have been demonstrated to be methane oxidizers from marine environments (82). These may be sediment associated or may recruit to the host to utilize algal products. The potential sulfate reducers within Desulfocapaceae may similarly recruit to the bell and its access to sediment pore waters (83). While *Pseudomonas* cf. *stuzeri* was not as high abundance as the other metabolizers listed above, it may have a role in denitrification or coral health for some species (84–86)

In addition to microbes with known metabolic functions that align with the *Cassiopea* holobiont, the genus *Ruegeria* includes many marine species and has been identified in the microbiomes of photsymbiotic and nonphotosymbiotic cnidarians alike (9,73,87). As Dirichlet modeling for bell microbiomes gave similar weight to many bacteria, the relative impact of all of these bacterial epibionts on the host is unclear.

### Laboratory cultures

*Cassiopea* are primarily used for in-lab studies on symbiosis (12,14,88). Some of these use wild-caught individuals, and others use lab-bred cultures, many of which have been held for years. The lab-raised *Cassiopea* used in Rothig et al. 2021 appear to have a completely different microbiome than the wild individuals caught here (19). This potential discrepancy should be considered when using wild-caught or captive-bred individuals, as captive-bred symbiosis and tolerance levels may differ from medusae in the wild. These discrepancies in microbiome are found in other model cnidarians, such as *Exaiptasia* (1). The information on wild *Cassiopea* microbial communities from the Florida Keys presented here may allow researchers working with lab-raised individuals to better contextualize the taxa that they find in cultured medusae.

## Conclusion

The microbiome of *Cassiopea* within the Florida Keys demonstrates strong body compartment discrimination and includes a central group of ASVs overrepresented in the gut and bell mucus. All of these families, Pseudomonadales, Vibrionaceae, and Mycoplasmataceae, are found in other medusae and anthozoans. Despite this overrepresentation of some microbial taxa within the gut, the *Cassiopea* bell is highly diverse, and overlaps with the communities of the water and substrate in and on which it resides. The consistency of *Cassiopea* microbial communities across the Florida Keys demonstrates that specific taxa are strongly associated with *Cassiopea* and that this can extend across many sites with diverse physical conditions. This work provides a greater understanding of the wild microbiome of the common lab organism *Cassiopea*.

## Acknowledgements

The present work was part of KMM’s PhD dissertation. Thank you to Dr. David Retchless for input on figure design. We thank the Florida Fish and Wildlife Conservation Commission for their help and guidance.

## Supplementary Information

Supplement A1: Full length 16S rRNA gene identities

Three samples were sent to CD Genomics for full length 16S on PacBio Sequel using universal 16S primers (F: AGRGTTTGATYNTGGCTCAG; R:TASGGHTACCTTGTTASGACTT). Samples sent for full length 16S rRNA sequencing included gastrovascular cavity swab extractions from medusae Marathon Key GVC 30 and Key West GVC 96 and GVC 120. As the usable reads returned from this were low (597-12520 per sample), these full-length identities are not discussed in the main text. For further details, please refer to the appendix (A1).

Full length 16S rRNA gene sequences (BioProject: PRJNA1020446) were run on mothur v1.48.0 with deviation from the MiSeq SOP on trim length. ASV cutoff was set to five differences, and ASV representative sequences were compared against the NCBI GenBank (Sept 24, 2023) database using BLAST to find nearest matches by e-value.

In order to identify core microbial groups to a lower taxonomic level, three internal medusa gastrovascular swabs (medusae Kmuffet_96, Kmuffet_120 and Kmuffet_30) from Marathon Key (MK) and Key West (GB) were sequence for full length 16S rRNA gene (see S1 for location of samples). As the primers were not chloroplast exclusionary, the Key West sites produced few useable sequences (Kmuffet_96: 597 and Kmuffet_120: 1097). The Marathon Key medusa (Kmuffet_30), likely bleached at time of collection, returned 12,520 non-chloroplast sequences. The Mycoplasma-like full length 16S rRNA gene sequence (Genbank Acc. No. OR592270) from Marathon Key and Key West has no closely related sequences currently identified (greatest identity 88.43% to sequence HQ393440). The *Endozoicomonas* strain from both Marathon Key and Key West (Genbank Acc. No. OR592271) has 99.93% identity to *Endozoicomonas atrinae* strain WP70(T) collected from the intestine of South Korean *Atrina pectinata* (Accession NR_134024)(89) and 99.80% identity (<5 nucleotide differences) with isolates collected from *Gorgonia ventalina* (GenBank Acc. No. GU118345) in Bocas del Toro, Panama (90). *Endozoicomanas* and *Mycoplasma-*like isolates were the only two groups to have ASVs with over 100 sequences.

S1 Tables: (in Supplementary Excel Workbook)

Supplementary Table S1: Roster of collected medusae

Supplementary Table S2: Read processing tracking table

Supplementary Table S3: Representative sequences of all ASVs from the dataset

Supplementary Table S4: ASVs orderd by abundance in each sample type

Supplementary Table S5: Taxonomy of top 1000 ASVs in dataset

Sfig 1: Rarefaction curves of all samples

